# Learning to read the imprints of consciousness on global brain dynamics: an application to intra-operative monitoring of anesthesia

**DOI:** 10.1101/225813

**Authors:** Leandro M. Alonso, Guillermo Solovey, Toru Yanagawa, Alex Proekt, Guillermo A. Cecchi, Marcelo O. Magnasco

## Abstract

In daily life, in the operating room and in the laboratory, the operational way to assess wakefulness and consciousness is through responsiveness. A number of studies suggest that the awake, conscious state is not the default behavior of an assembly of neurons, but rather a very special state of activity that has to be actively maintained and curated to support its functional properties. Thus responsiveness is a feature that requires active maintenance, such as a homeostatic mechanism to balance excitation and inhibition. In this work we developed a method for monitoring such maintenance processes, focusing on a specific signature of their behavior derived from the theory of dynamical systems: stability analysis of dynamical modes. When such mechanisms are at work, their modes of activity are at marginal stability, neither damped (stable) nor exponentially growing (unstable) but rather hovering in between. We have previously shown that, conversely, under induction of anesthesia those modes become more stable and thus less responsive, then reversed upon emergence to wakefulness. We take advantage of this effect to build a single-trial classifier which detects whether a subject is awake or unconscious achieving high performance. We show that our approach can be developed into a mean for intra-operative monitoring of the depth of anesthesia, an application of fundamental importance to modern clinical practice.

## I. INTRODUCTION

General anesthesia is a common medical procedure by which a human patient becomes unconscious to the point that invasive surgeries can be performed without inflicting pain or awareness of the procedure. But how can we be sure that no pain is being experienced or that the patient is not aware? In practice, this delicate task is ultimately left to the expertise of the anesthesiologist. Despite vast accumulated experience on these practices, the anesthetic procedure is still imperfect and a small but significant fraction of patients experience anesthetic awareness^1^. There is still to date an alarming incidence of anesthetic awareness, whereby patients awaken during a major surgical procedure, often unable to communicate their predicament. The reason this continues to occur is simple: we currently have no means of differentiating the brain activity of conscious patients from that of anesthetized patients. To put it another way, the central difficulty is the lack of any operational way of quantifying pain or consciousness. A major complication is that the large number of anesthetic agents in routine surgical use act on different pathways, involve different loci, and cause different changes in electrical activity.

Changes in the level of arousal (wakefulness) have been historically quantified using spectral analysis of neuronal activity. In this view, decrease in the level of wakefulness is reflected in the increase and prevalence of low frequency oscillations and the concurrent decrease in the high frequency oscillations^2^. While this is true for some states of decreased arousal such as slow wave sleep, this association breaks down during other states in which arousal is similarly depressed such as rapid eye movement (REM) sleep for instance. Furthermore, state of general anesthesia can be characterized by different spectral signatures depending on the specific choice of anesthetic agent^3^. This makes current modes of detecting the “depth of anesthesia” unreliable^4^.

It has been suggested that neural systems operate in a critical regime similar to phase transitions in physics, given several computational desirable features of such states represented by the statistics of the thermodynamic variables^5^. Evidence for *statistical* criticality is based on the observation that various aspects of neuronal activity such as avalanches observed in local field potentials and action potentials in tissue preparations and in animal models^6,7^, as well as magneto-encephalography (MEG) and electro-corticography (ECoG) in human subjects^8,9^, exhibit long tailed-distributions well approximated by power laws. More recently, the *dynamical* aspect of criticality has been brought into focus, as a similarly desirable feature not fully captured by steady-state statistics such as avalanche size distributions^10–12^; a perturbation in an extended dynamical system that is close to a critical point will neither decay nor explode, thus allowing for long range communication across the entire system. In contrast, if the system is far from criticality (therefore stable), perturbations damp out and no information integration takes place beyond the characteristic time scale which characterize the damping. The critical regime provides important functional benefits; quantities such as dynamic range and information transmission are optimized near criticality^13^. If indeed dynamical criticality is a useful feature of brain activity, stability of neuronal dynamics ought to be modulated by the behavioral state of the subject. When the brain is awake and displaying complex statistical behavior its dynamical state ought to be close to a bifurcation point; marginally stable modes contribute to long range interactions across the system. Conversely when higher-order functions associated with wakefulness have been diminished and eventually completely shut down by anesthesia, brain dynamics should exhibit more stability. In other words, anesthesia induction should lead to stabilization of brain dynamics.

To address these questions, we fitted vector autoregressive (VAR) models to electrocorticography recordings (ECoG). These are routine measurements which are performed in human subjects with chronic epilepsies. A regular grid of electrodes is placed on the surface of the exposed brain and electrical activity is recorded. These measurements correspond to the synchronized activities of thousands of neurons and offer a unique window to cortex brain activity. Our previous results suggest that tracking changes in the dynamical stability of (VAR) models fitted to these signals yields a marker which covaries with the level of arousal of the subjects: the dynamical global modes of brain activity, which under normal circumstances hover between stability and instability, become more stable^17,18^. We have demonstrated that this specific hallmark is disrupted in anesthesia and restored after recovery, in humans and macaques, for two different anesthetic agents^17,18^. A number of studies suggest that monitoring the departure from critical dynamics may provide a useful neural correlate of conscious behavior^19,20^. ECoG recordings provide the basis for several studies of conscious function^21^. Here we study ECoG recordings collected directly from the the full cortex of non-human primates as they were gradually induced into the state of general anesthesia.

Our previous studies have shown statistically significant differences in stability for each specific individual by aggregating a fair amount of data. However this does not permit looking at a few seconds of activity from one patient and state whether the patient is awake or unconscious, because the baseline for each individual is unknown. In this work we develop a single-trial classifier which aims to determine the state of the subjects using short temporal snippets of the ECoG recordings. We build vectors based on the dynamical stability (DS) of the ECoG signals and train support vector machines (SVM) to classify the subjects states. We test our procedure in two situations of clinical relevance achieving high performances. Our results suggest that one may be able to build a baseline-free classifier using measures based on dynamical stability.

## II. METHODS

### Subjects and data acquisition

Data from four male monkeys were collected at the Laboratory for Adaptive Intelligence, Brain Science Institute, RIKEN. Electrocorticographic (ECoG) recordings were sampled at 1 kHz from an array consisting of *N* = 128 electrodes covering both full hemispheres. A more detailed description of the experiments can be found in [Nagasaka et al., 2011; Yanagawa et al., 2013]. ECoG recordings were obtained during the induction of anesthesia starting from the awake state. The dataset consists also of video footages of the experiments in which behavioral assessments are performed to determine loss of consciousness. This dataset is not available to the public. In this study we analyze a total of 16 experiments each consisting of reversible induction of anesthesia starting from the awake state.

### Description of the dataset

A total of 12 anesthetic inductions were performed using ketamine medetomidine (KM) doses. Four anesthetic inductions were performed with propofol (P). Ketamine medetomidine inductions were performed by injecting the drugs intramuscularly, whereas propofol was administered intravenously. Each monkey received more than one anesthetic induction that were separated by at least 1 day. We labeled our subjects as (M1,M2,M3,M4). Our dataset consists of 4 sessions of KM for M1, 2 sessions of P for M1, 3 sessions of KM for M2, 3 sessions of KM for M3, 2 sessions of KM for M4 and 2 sessions of P for M4.

### Data processing

All channels were notch filtered to eliminate electrical line noise at 50*Hz*, 100*Hz* and 150*Hz*. Then we applied a bandpass filter between 5Hz and 500Hz. Both notch and bandpass filters were implemented using the idealfilter function in MATLAB (MathWorks) to avoid phase shifts. These procedures were also performed using the python scipy function filtfilt yielding identical results.

### Stability analysis

The notion that the brain might be operating in a critical regime has been explored by many authors. Dynamical systems theory indicates that systems which are capable of performing computations should have a large number of modes with marginal stability. In such a scenario an arbitrary perturbation wont decay nor explode, thus allowing for information integration across the entire system. Therefore, it has been suggested that the brain might operate in a dynamically critical regime. A simple model exhibiting complex spatio-temporal dynamics was recently proposed by Magnasco et al. in which statistically critical behavior emerges due to dynamical instabilities^10^. Overall, theoretical considerations suggest that when the brain is awake its dynamical state is close to dynamical criticality, a state in which many dynamical degrees of freedom are neither unstable (they do not explode) nor stable (they do not decay) but straddling the interface between stability and instability. These marginally stable modes contribute to long range interactions across the brain. This leads us to associate wakefulness to dynamical marginality; conversely, the anesthetized brain should exhibit more damping in those modes that are associated to conscious function or cognition. If this view is correct, a measure of the dynamical stability of the system could then be used as a marker for depth of anesthesia^17^.

ECoG is a multivariate time series whose dynamical properties can be inferred using autoregressive models fitted independently to short time segments as described previously^24^. In order to test the notion that consciousness can be associated to a dynamical homeostatic mechanism, we assume locally linear dynamics in short temporal snipets of the recordings and fit vector autoregressive (VAR) models. This allows us to address changes in the stability properties of the fitted linear approximation as the concentration of anesthetics is increased. For a given temporal interval of duration *δ* we fit an order 1 VAR model. This is the simplest linear dynamical system that can be fit to a multivariate time series.

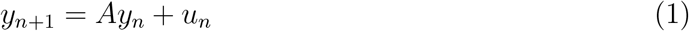

Here, *y_n_* ∈ ℜ^*N*^ is a multivariate time series that corresponds to the recorded activity in all *N* =128 channels at time *t_n_, A* ∈ ℝ^*N×N*^ is the matrix to be estimated and *u_n_* is assumed to be white noise. A comprehensive treatment of this model and its estimation can be found in Lütkephol^29^.

In this work we used a statistical modeling module for python called Statsmodels^36^. The idea behind the estimation procedure is that for a temporal segment containing n datapoints, one ends up with a system of *n* linear equations of the form 1. Then finding A under the assumption of uncorrelated noise can be cast a estimating the pseudoinverse of the matrix containing the *y_n_* values. This is in turn performed by the standard python library numpy lstsq module^38^. Finally, this module is a python interface for the popular package LAPACK which contains several algorithms for performing singular value decompositions of large matrices^37^. The performance of our procedure relies ultimately on this standard and heavily tested library.

Each time a model is fit to data we obtain a matrix A which governs the stability properties of the VAR model. In order to address changes in the dynamical stability of the fitted models we consider the distribution of eigenvalues of A. Since our underlying hypothesis corresponds to a continuum model we performed a transformation in order to obtain a correspondence between the eigenvalues of A and the timescales of the dynamics. Let *λ_j_* = *ρ_j_e^iϕ^* be the eigenvalue corresponding to the *j*-th mode, the frequency of the mode is given by 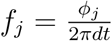 while the growth rate (timescale) of the mode is given by 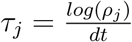. Here 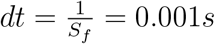, where *S_f_* = 1000*Hz* is the sampling frequency of the recordings.

Our previous results suggest that as the subjects become anesthetized the linear stability of the ECoG recordings exhibits significant stabilization which is efficiently quantified by non parametric statistical methods. Such stabilization effect was first reported in recordings performed in human patients with temporal lobe epilepsy while being anesthetized with propofol [Alonso et. al. 2014^17^]. This finding was further supported by performing this analysis on the current dataset in monkeys [Solovey & Alonso 2015^18^]. This led us to associate loss of consciousness with stabilization of cortical activity. In this work we explore the possibility that such stabilization effect can be exploited to predict conscious activity.

### DS vectors

Dynamical stability (DS) is determined by the fitted distributions of eigenvalues. For a given temporal segment of duration Δ ≥ 500*msecs*, we build (DS) vectors based on this measure by coarse graining this distributions. We sample the distributions by fitting 10 equally spaced VAR(1) models using *δ* = 500*msecs* windows independently. Note that for choices of Δ ≤ 5000*msecs* the models fit overlapping data. Vectors are obtained by binning the distributions of eigenvalues in a 20 × 20 grid in the range 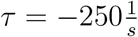 to 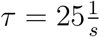 and *f* = 4*Hz* to *f* = 256*Hz*. The frequency axis is in logarithmic scale base 2. The range and scales were chosen to highlight the differences between the awake distributions versus the anesthetized ones. About half of the total eigenvalues fall within this range. The vectors are obtained by flattening the distributions; this is horizontally stacking the row values of the grids yielding a vector with dimension 400.

### FFT vectors

In order to compare (DS) performance against a spectral method we build FFT vectors based only in spectral features of the data. For each channel, we compute the fast fourier transform of the data contained in an interval of duration Δ and keep the logarithm of the power *p* for frequencies smaller than 100Hz; *υ* = *log*(*p_f<100Hz_*). A vector is defined as the average of *p* across channels. The dimension of the resulting vector then depends on the duration of the interval Δ. Performance is largely independent of this dimension.

### Data labeling

Each vector was labeled as awake or anesthetized by careful assessment of the experiment videos. For each experiment two temporal intervals were determined. The awake interval corresponds to resting with eyes closed condition, prior to any drug injection. The anesthetized intervals occur always after drug injection and were determined as the interval in between two behavioral assessments performed by the experimenters in which the subjects did not respond. Responsiveness assessments were in most cases tactile and in some cases the subjects were also perturbed with noise. Both intervals were further shortened by removing the first and last 30 seconds to decrease the chance of mislabeling. The interval durations are in general different across experiments ranging from 350*secs* to 1500*secs* for Ketamine Medetomidine. The anesthetized intervals are much shorter in the case of propofol ranging 130*secs* to 450*secs*. In all experiments the awake and anesthetized intervals are separated by at least 400*secs*.

### Surrogate tests

We applied three surrogate procedures to the ECoG recordings. Phase surrogation is a standard procedure which consists of Fourier transforming the data, randomizing the phases and transforming back. We applied this surrogation to each channel and used the surrogated data to build vectors. Staggering surrogation corresponds to desynchronizing the data locally. Each channel is shifted forward in time by a random value taken from a flat distribution of width 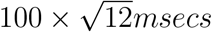. This is performed every time an interval is selected for building a vector. Global staggering surrogation corresponds to the same procedure except that the shifts are determined only once and the same shifts are applied to all vectors of the same experiment. Global staggering surrogation was performed by taking the lags from a flat distribution of width 500*msecs*. All surrogate tests aim to disrupt global information across recording sites.

### Classification method

For each experiment we built a dataset by taking 500 equally spaced vectors on the awake interval and 500 equally spaced vectors in the anesthetized interval. For each dataset we also generated a surrogate dataset consisting of vectors built with surrogated data. The same procedure was also applied to obtain datasets for varying amounts of data Δ in order to test for performance as this parameter is varied. Depending on the choice of this parameter and the duration of the awake and anesthesia intervals of each experiment, adjacent vectors within an interval may be constructed with partially overlapping data.

We used support vector machines (SVM) for classification. SVMs in their simplest form are linear classifiers which are particularly efficient in high dimensional spaces and for which there are efficient training methods. In this work we implemented the machine learning python library scikit-learn^35^ to train SVMs with linear kernels for classifying whether a subject is awake or unconscious.

### Training and testing protocol

We test our classifiers on unseen data. Given a train dataset (T) and test dataset (P) to assess performance we proceed as follows. We take half the vectors on (T) *at random* to train the SVM and then ask the classifier to predict the full (P) dataset of a different subject. We then define the error as the number of wrong classifications in percent. Thus, one source of variability in the errors when this protocol is repeated comes from different choices for the training vectors.

## III. RESULTS

A total of 16 experiments were analyzed in this work. Figure 1A shows the electrode placement and Figure 1B shows the recordings of channels 1 through 32 (indicated by red dashed lines in Fig. 1A) for a short temporal segment (500*msecs*). As a way to track changes in the dynamical stability (DS) of the fitted VAR(1) models as anesthesia is induced, we compare the initial distribution of damping time scales (*Re(λ)*) against subsequent distributions using a Kolmogorv-Smirnov test. Figure 1C shows the KS value of the test along the course of a full experiment in one subject, for each anesthetic. First (*t* < 1000*secs*), the subjects are resting with their eyes open and no drugs are given. At about *t* = 1000 the subjects are blindfolded and this changes the stability of the models. Drugs are given at time *t* ≈ 2200*secs* and responsiveness assements are performed to determine the state of the subject. The subjects recover from anesthesia and the blind is removed (*t* > 5500). For each experiment we determined the intervals of the awake state and the anesthetized state by assessing the experiments videos.

**FIG. 1.**
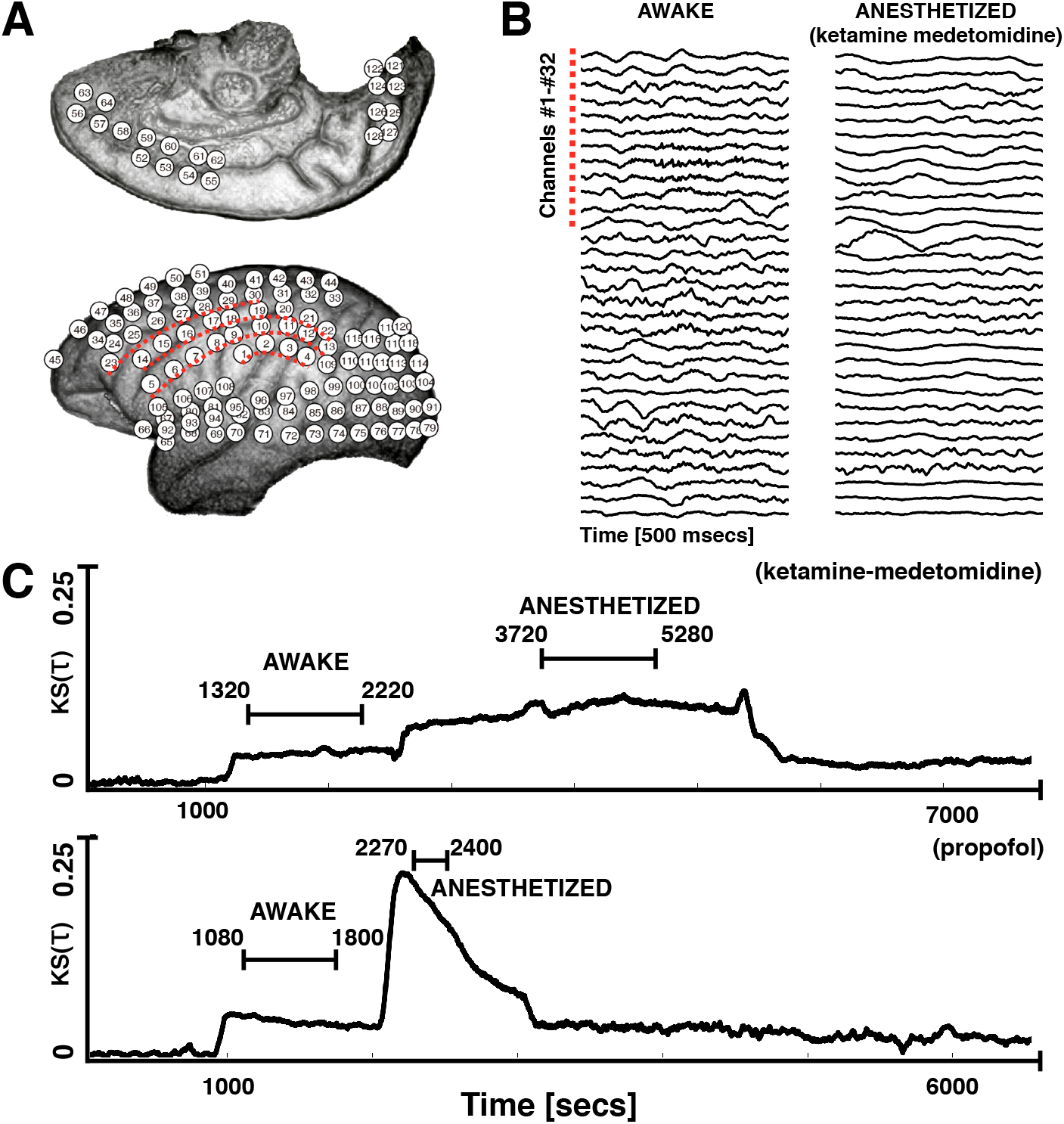
Changes in dynamical stability as subjects are induced into anesthesia. We computed the dynamical stability of VAR(1) models fitted to the ECoG recordings as the subjects underwent general anesthesia. The intervals labeled as *awake* and *anesthetized* were determined from responsiveness assesments performed during the experiments. **A** Electrode placement **B** Data recorded from channels 1 through 32 for the awake condition (left) and the anesthetized condition (right) undeter ketamine-medetomedine. **C** We quantified changes in the stability of the models by comparing the distributions of damping time scales (*Reλ*) using a Kolmogorov-Smirnov test. The y-axis corresponds to the KS coefficient of comparing the initial reference distribution (awake, no drugs) versus the distributions of damping time scales obtained at subsequent time stamps.

Our study aims to determine whether DS can be used to predict whether a subject is conscious or not by using a short temporal segment Δ of the ECoG recordings. For this we imagine two situations of clinical relevance. First, the awake and anesthetized states for three subjects are known and this information is used to train a classifier. The trained classifier is then used to predict the state of the subject left out for every session of the same drug (KM). The second scenario we tested is when the awake and anesthetized activity for an individual is known and the outcome of the next session for the same drug is predicted. We computed the dynamical stability (DS) of the models fitted to the signals in the awake and anesthetized intervals and obtained vectors from binning the distributions of eigenvalues (see methods). We then trained support vector machines (SVM) classifiers on subsets of the experiments and tested the resulting classifiers on the remnant unseen subsets. We compare the performance of our protocol (DS) with spectral measures (FFT) and DS vectors built with surrogate data. Throughout this work, the classifiers are tested on unseen data.

Figure 2A shows the vectors obtained for subject M1 for both drugs and for both the awake and anesthetized conditions. The distribution of eigenvalues of the models fitted to the recordings is binned in a range and the vectors are built by keeping the count in each bin. The awake and anesthetized distributions exhibit remarkable differences for both anesthetic agents. These differences are highlighted in the overlay plots in row (Fig. 2B). The awake distribution is represented in cyan while the anesthetized distribution is represented in red. The points in which these distributions have similar occupancy are then represented in white. Our previous work suggests that loss of consciousness is concomitant with increased stability of the fitted models (more negative damping timescales *τ*). This observation is consistent with the plots shown in Fig. 2C. The 2D distributions of eigenvalues for both states are integrated along the frequency axis to obtain the distributions of damping timescales *τ* for the awake (blue) condition and the anesthetized (red) condition. We performed the same analysis using phase surrogates of the data (see methods). Our surrogation procedure aims at disrupting correlations amongst channels that may contain global information. We find that by performing these disruptions the dynamical stability of the models changes noticeably while still exhibiting consistent differences across states and drugs. These analysis are shown in rows B and C and it is indicated in the figure labels.

**FIG. 2.**
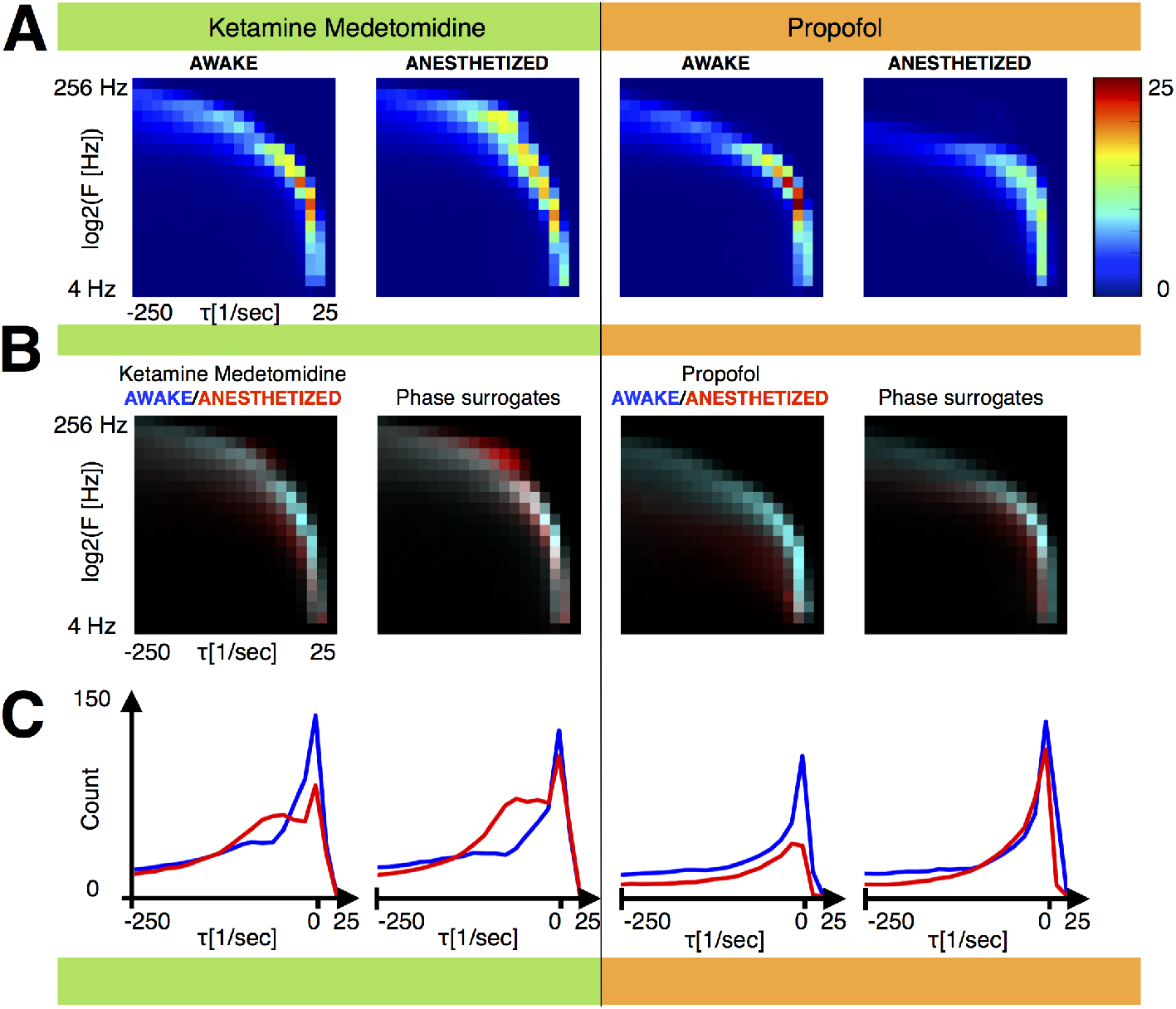
Description of the dataset. We used the distributions of eigenvalues of the fitted VAR models to build vectors by computing a 2D histogram over a range. Mean vectors for M1. Left panels: Ketamine Medetomidine. Right panels: Propofol. **(A)** Mean DS vectors for awake and anesthetized conditions. **(B)** Overlay between awake (blue) and anesthetized (red) vectors. Histograms were normalized between 0 and 1 and the overlay highlights changes in the distributions under both conditions. For each drug we plot the mean DS vectors overlay and the mean DS vectors using surrogated data (phase randomization). **(C)** Distribution of damping timescales *τ* for awake (blue) and anesthetized (red) conditions, for real and surrogate data. The vectors were built using Δ = 2000*msecs* of data.

Figure 3 shows the results of training the classifiers on three subjects and predicting all the experiments for the subject left out. In this case the training (T) set corresponds to stacking all the vectors for the KM condition in three subjects and the test set (P) is obtained by stacking all the vectors for the KM condition in the subject left out. The vectors were built using Δ = 2*secs* of data (see methods). The figure corresponds to the histograms of scores obtained for *N_f_* = 1000 folds. In order to obtain a measure of the statistical power of the classifier, we computed the ROC curve for the best classifier. This curve computes the tradeoff between false positives and false negatives as the discrimination threshold of the classifier is varied. The area under the curve (AUC) is a measure of the statistical power of the classifier and we indicated it in the figure labels.

**FIG. 3.**
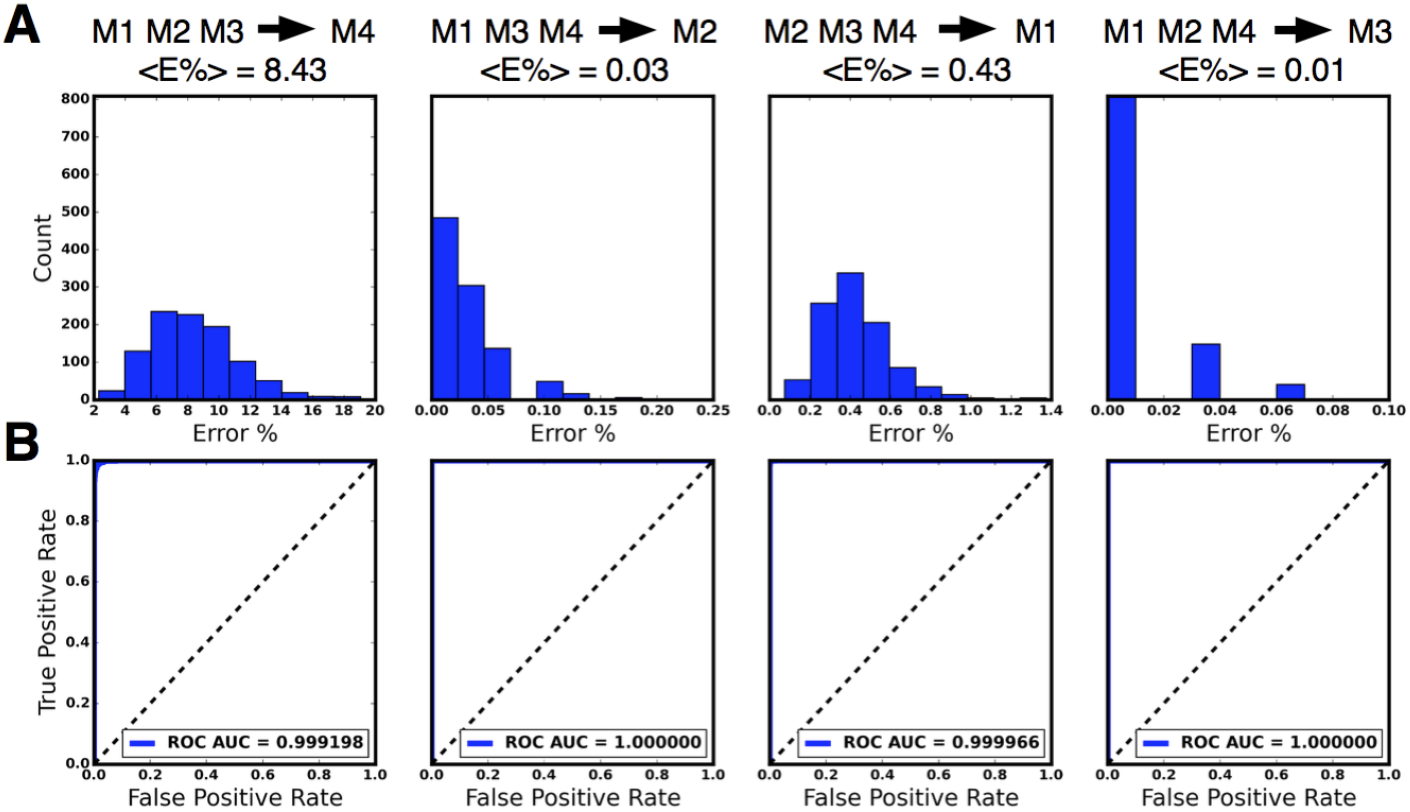
Best case performance across subjects (Ketamine Medetomidine). We test the performance of the protocol using datasets for three subjects for training the classifiers and used the subject left out for testing. We chose a window size Δ = 2000*msecs* of data to build the vectors. **(A)** Histograms show the error of the classifiers for *N* = 1000 folds of the protocol. **(B)** Receiver operating characteristic (ROC) curves showing the performance of the best classifier as its discrimination threshold is varied. The area under the curve (AUC) is indicated in the figure labels.

**FIG. 4.**
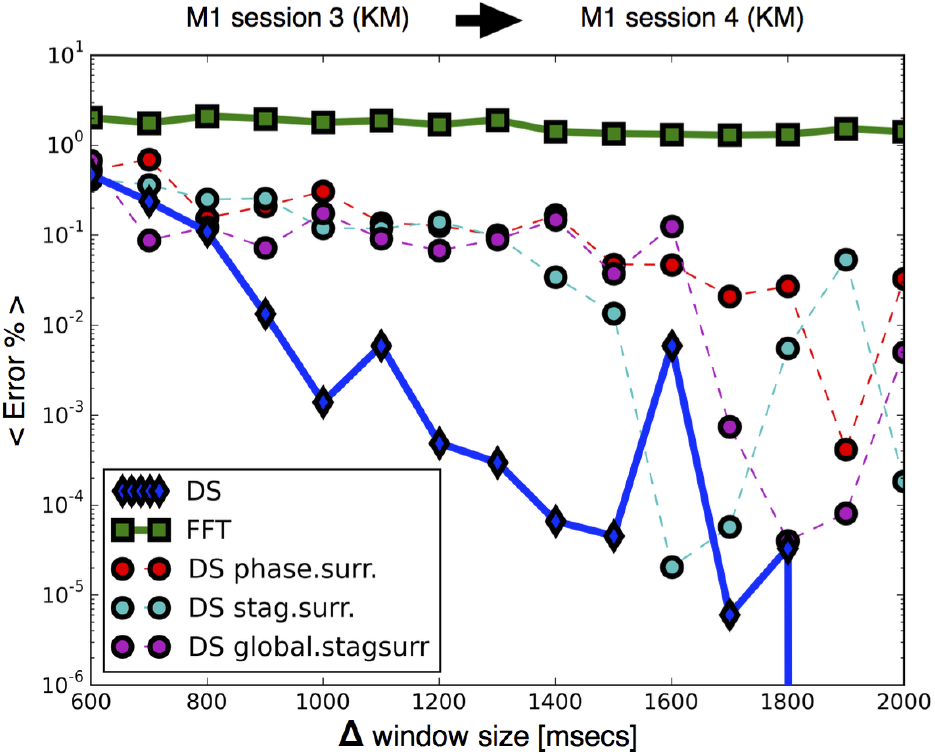
Performance increases as more data is included in the vectors. Improvement in performance for increasing window size Δ across sessions (Ketamine Medetomidine): we test the performance of the protocol for increasing amounts of data that is used to build each vector. We train the classifiers in M1 session 3, and predict the state of the same subject M1 in session 4. The plot compares the performance of the task on vectors built with surrogate data against vectors built with undisrupted data. Dynamical stability (DS) outperforms a local spectral method (FFT) and disruption of global information leads to worse performances. The number of folds at each point is *N_f_* > 10^5^

Figures 4 and 5 show the performance of the protocol as more data is utilized to build the Ketamine Medetomidine vectors. We performed this test by training the classifiers with vectors from one session of M1 for KM induction (T), and then tested our predictions on a dataset corresponding to the same subject M1 and the same anesthetic KM, for a different session (P). The figure is comparing the performance of our protocol (DS) against different datasets. Dataset (FFT) is obtained based solely on spectral methods while the surrogates correspond to DS vectors built using surrogated data. The figure shows that (DS) outperforms (FFT) and interestingly, if data is disrupted the performance is worsened by roughly two orders of magnitude. This suggests that our method is picking global dynamical features of brain activity and that these features are useful to predict consciousness and lack of it. It is important to note in this case that while (DS) outperforms (FFT) this also happens for (DS) vectors built with surrogate data. Our surrogation tests show that bulk stabilization of the distributions under anesthesia is moderately resistant to these disruptions. When vectors are built with surrogated data, the distribution of eigenvalues still exhibits consistent differences across states. This is in turn consistent with comparable performances of the classification tasks on real and surrogate data vectors.

**FIG. 5.**
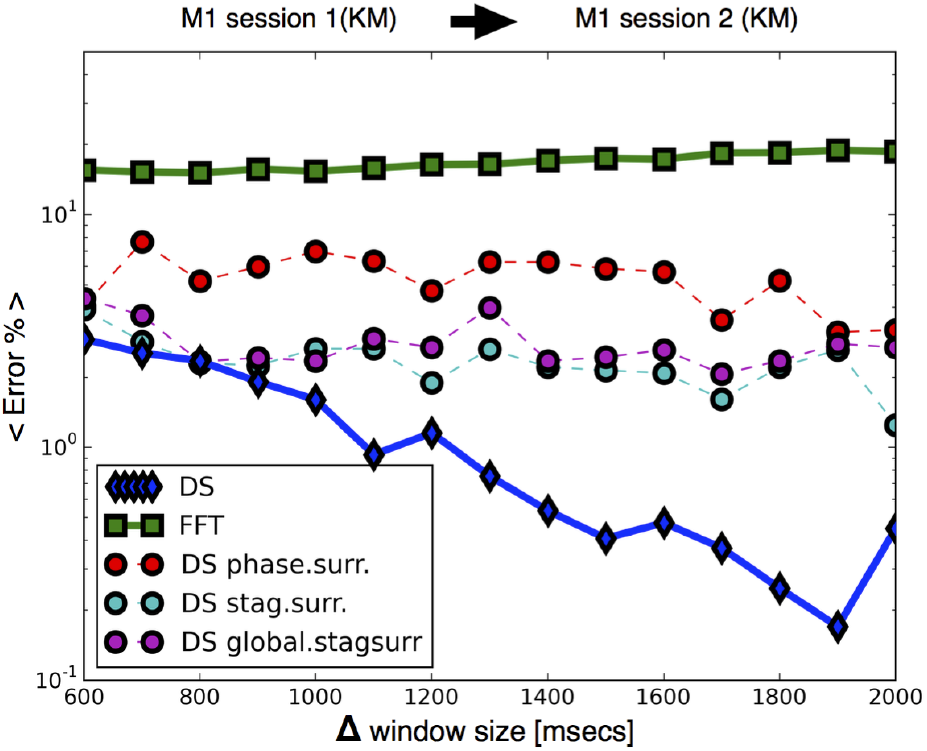
Improvement in performance for increasing window size Δ across sessions (Ketamine Medetomidine): we test the performance of the protocol for increasing amounts of data that is used to build each vector. We train the classifiers in Ml session 1, and predict the state of the same subject M1 in session 2. The number of folds at each point is *N_f_* > 10^4^

In Figure 6 we show the same analysis as in Figures 4 and 5 for inductions with Propofol across sessions for subject *M*1. In this case, the performance of the classifiers on the (DS) vectors is about 3 to 4 orders of magnitude better than on the (FFT) vectors. Interestingly, in this case the (FFT) vectors perform similarly to the (DS) vectors built with surrogate data. This is also consistent with Figure 1 row C in which we show that while in the Ketamine Medetomidine case, bulk differences across states persist after surrogation, for the case of Propofol differences across states depend more strongly on global correlations which are disrupted by surrogation. Performance across subjects for propofol is not presented in this work since our analysis suggests that M4 does not become fully anesthetized with this drug. This in turn could be due to the fact that M4 belongs to a different species than the other subjects.

**FIG. 6.**
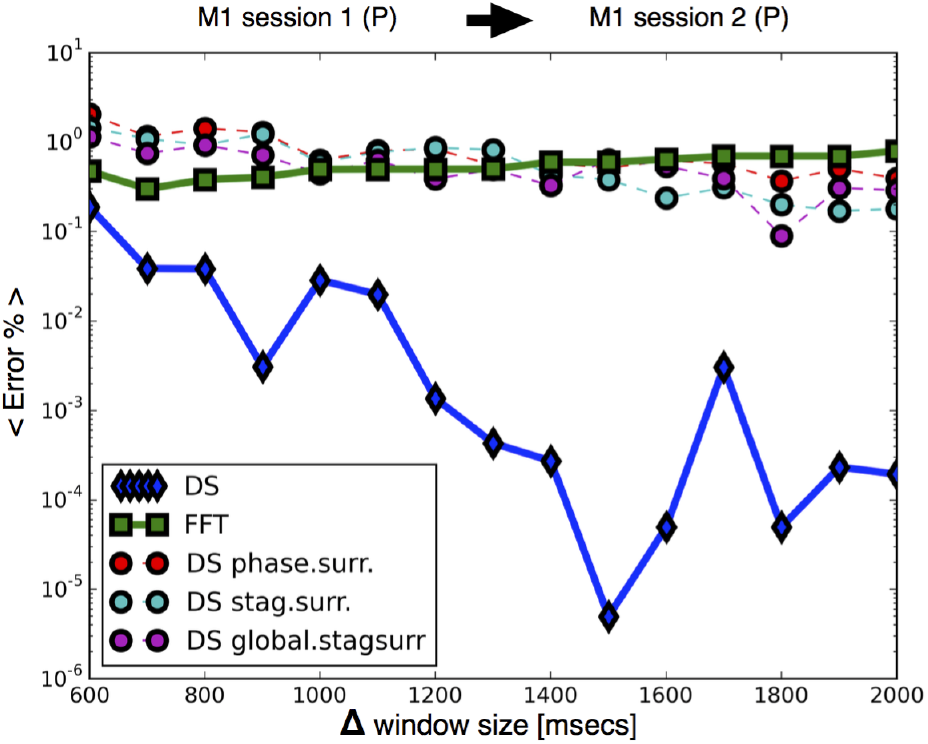
Improvement in performance for increasing window size Δ across sessions (Propofol). Dynamical stability (DS) outperforms a local spectral method (FFT) and disruption of global information leads to worse performances which are similar to (FFT). The number of folds at each point is *N_f_* > 10^4^

We found that the performance on real data is larger than on surrogate data for both anesthetics. This suggests the exciting possibility that the classifier is picking up on global dynamical features which are disrupted by surrogation. Our current results do not demonstrate that the global feature that is being picked up is similar across monkeys and across drugs. However, our results suggest that a simple linear approach is able to capture global features that are associated to loss of consciousness. Our numerical estimates provide a measure of the utility of such global information. Consistent with the fact that the stability of the fitted models is to a certain degree independent of global correlations, the performance of our procedure is only moderately worsened by disruption of such global features. This suggests that high performances might be achieved by these means even in the more common situation in which a smaller portion of the cortex is being monitored

## IV CONCLUSIONS

Unveiling the mechanisms by which consciousness emerges is among the ultimate goals of systems neuroscience. Within this broad scope, a more immediate goal is understanding the quantitative imprint of consciousness on electrophysiological activity. Our efforts aimed at establishing a relationship between brain activity under different anesthetic regimes and the linear stability of the registered ECoG signals. Our previous studies have shown that under induction of anesthesia the dynamical stability (DS) of linear models fitted to the ECoG signals is increased and thus become less responsive, then reversed upon emergence to wakefulness^17,18^.

We have developed a single-trial classifier based on this measure using support vector machines. Our results indicate that dynamical stability (DS) can be used to determine whether a subject is conscious or not with remarkable accuracy. We also found that by performing these procedures we outperform other single-channel measures based only on spectral properties (FFT). According to the literature this procedure has the potential to outperform other previously published procedures.

Our method is novel since it aims to fit global brain dynamics as opposed to relying on many single-channel measurements. The assumptions behind our procedure are minimal, namely, that the dynamics is linear in short temporal intervals. This assumption enables efficient estimation procedures and we kept the simplest approach within this premise. Our results suggest that it is likely that a global feature which could be associated to consciousness may be revealed by a many-channel linear approach. Interestingly, we found that the classifiers are using global information to increase performance to a certain degree. This is exciting because if the presence of a global dynamical feature is important to increase performance in a consciousness detection task, there are chances this can be linked to consciousness provided we can show consistency across anesthetics and subjects. Our current approach does not probe the possibility that the global feature that is being picked up is similar across monkeys and across drugs. On the other hand, our approach demonstrates that fitting the simplest dynamical system to the data has the potential to outperform other methods to monitor for loss of consciousness.

We found a potentially important clinical application. Our procedure yields unprecedented performance in several consciousness detection tasks of clinical relevance. The procedure makes use of information on the global dynamical state of the brain to further increase performance. It cannot be discarded that such global features can be linked to consciousness and additional tests with other anesthetics are required to establish a stronger connection. Consistent with the fact that the stability of the fitted models is only moderately dependent on correlations, the performance of the method is also moderately worsened by disruption of such global features. On the other hand, we showed that if global information is not disrupted, then the procedure consistently performs better. We believe that this procedure has the potential to outperform other methods currently utilized to monitor for depth anesthesia. Further tests with other anesthetics are required to attempt a thorough translational effort. Extensions of these methods for non-invasive recordings as EEG as well as invasive recordings as Utah arrays is highly desirable.

## ACKNOWLEDGMENTS

LM Alonso’s research was supported by funds from a Leon Levy Fellowship at The Rockefeller University.

